# Software application profile: GLU: A tool for analysing continuously measured glucose in epidemiology

**DOI:** 10.1101/500256

**Authors:** Louise A C Millard, Nashita Patel, Kate Tilling, Melanie Lewcock, Peter A Flach, Debbie A Lawlor

## Abstract

**Motivation:** Continuous glucose monitors (CGM) record interstitial glucose ‘continuously’, producing a sequence of measurements for each participant (e.g. the average glucose every 5 minutes over several days, both day and night). To analyze these data, researchers tend to derive summary variables such as the Area Under the Curve (AUC), to then use in subsequent analyses. To date, a lack of consistency and transparency of precise definitions used for these summary variables has hindered interpretation, replication and comparison of results across studies. We present GLU, an open-source software package for deriving a consistent set of summary variables from CGM data.

**General features:** GLU performs quality control of each CGM sample (e.g. addressing missing data), derives a diverse set of summary variables (e.g. AUC, and proportion of time spent in hypo-, normo- and hyper-glycaemic levels) covering six broad domains, and outputs these (with quality control information) to the user.

**Implementation:** GLU is implemented in R.

**AVAILABILITY:** GLU is available on GitHub at [https://github.com/MRCIEU/GLU]. Git tag v0.1 corresponds to the version presented here.

## INTRODUCTION

Epidemiological and clinical studies interested in circulating glucose as a risk factor or outcome typically measure levels in the blood (fasting, non-fasting and/or post-oral glucose) at a single or widely spaced time points (e.g. every few years) (1–4). While these are important health indicators, there has been an increasing appreciation that glucose levels and variability in free-living conditions during both the day and night, may also provide important health measures in clinical (e.g. diabetic or obese) and ‘healthy’ populations (5–11). Continuous glucose monitoring (CGM) systems measure interstitial glucose levels by implanting a sensor subcutaneously (12). Typically, finger prick blood glucose measurements are needed to calibrate the interstitial glucose levels to capillary blood glucose levels, although devices that do not need this calibration step are now becoming increasingly available (12,13). Throughout this paper we refer to the sensor predicted capillary glucose levels as ‘sensor glucose’.

CGM systems were initially used in research evaluating their potential value in patients with diabetes, and are now increasingly used in the management of type I and type II diabetes (8,9,14–16). More recently, CGM has been used in a wider-range of epidemiological studies. For instance, CGM has been used to measure glucose levels ‘continuously’ over a number of days to identify hypo-glycaemia in those receiving intensive care, and in ‘healthy’ populations to explore whether it can be used to identify groups at increased risk of diabetes, including gestational diabetes (17–21). Unlike glucose assessed at a single time point providing only a ‘snap-shot’ of glycaemic control, or glycated haemoglobin that gives a single measure indicating mean glucose levels over a period of weeks, researchers can use these continuously measured data to assess how glucose levels vary across the day and night for several days or weeks and identify determinants of this variation and its health impact (17–21).

Researcher using CGM data tend to first derive summary variables that are then used in their subsequent analyses (e.g. exploring the association of these summary variables with later health outcomes). Summary variables might include area under the curve (AUC) (i.e. the average glucose level over time) or time spent in low, medium or high levels. While there are a set of variables that may be commonly derived in CGM studies there are increasing examples of studies addressing broadly similar research questions but deriving different summary variables. For example, we found two papers assessing glycaemic variability in non-diabetic people, one that included morbidly obese participants (17) and the other that included healthy people (22). Whilst both of these studies used standard deviation (SD), coefficient of variation (CV) and mean amplitude of glycaemic excursions (MAGE) as measures of variability, the one in morbidly obese people also used mean of daily differences (MODD) (17) and the other used mean absolute rate of change (MARC) (22). These two studies illustrate that (a) several measures of variability can be derived from CGM data and it is important to justify which are used and differences between them, which neither of these papers did, and (b) we would want consistent measures to be used across studies. Even when different studies derive a variable representing the same fundamental property it may be defined differently, for example using different thresholds to define hypo-, normo- and hyper-glycaemia (5,17). This lack of consistency across studies, together with insufficient reporting of study methods, means that it is difficult to interpret results. It is also difficult to seek replication or pool study results in meta-analyses, when varied measures are derived (5–8,11,23–25). For example, a recent review that compared studies according to the proportion of time in hypo-normo- and hyper-glycaemia was limited because researchers used different thresholds or did not include these measures at all (17). It is also unclear whether researchers derive many summary variables but only present those for which analysis supports their hypothesis, such that the evidence published in the literature and on which clinical decisions are based may be biased (26). The American Diabetes Association recently suggested some summary statistics (such as the coefficient of variation to assess variability and proportion of time in ranges [hypo-, normo- and hyper-glycaemia]) to assess glucose control in patients with diabetes but acknowledged further research was needed to establish which summary measures are most useful even in diabetes patients (8). Outside this guidance we are unaware of any that has been suggested for the broader use of CGM in epidemiology; nor are we aware of any general epidemiology research tools to systematize analyses of CGM data.

In this paper, we present GLU, a general open-source tool for processing CGM data. GLU performs quality control and derives a set of glucose characteristics (illustrated in Figure 1), that can be used in subsequent analyses. Use of a common tool will help to standardise methods across research studies. Hence, in the future it will be easier to compare and meta-analyse results across studies, and perform replication analyses. An open source tool also improves transparency of methods as all code is freely available, aiding interpretation of results. Furthermore, we intend to update GLU as methods advance. The presentation of this tool is timely as CGM is beginning to be widely adopted in epidemiological research, including both observational studies and randomised controlled trials (17–21).

**Figure 1:**
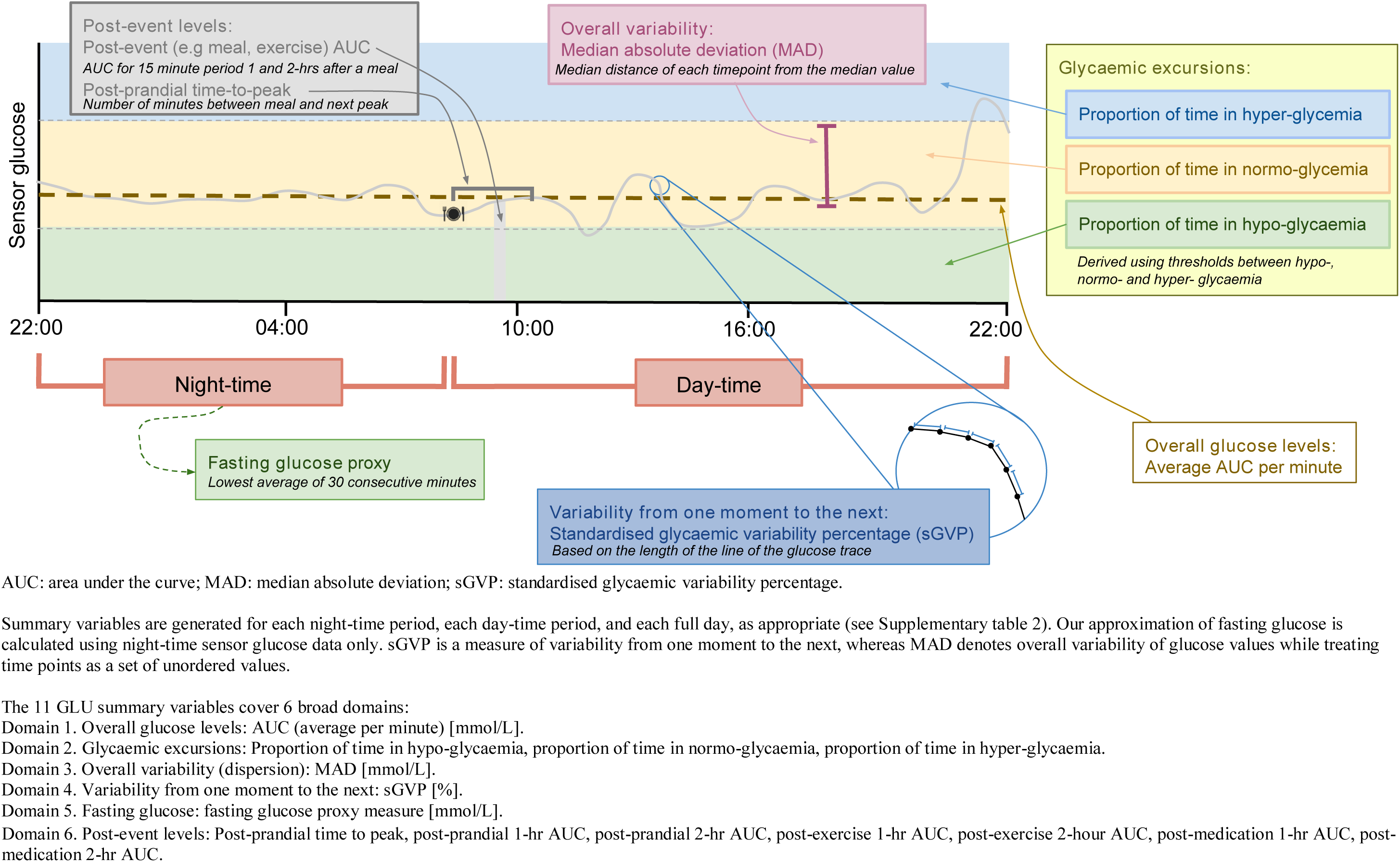
Illustration of summary variables derived by GLU

## IMPLEMENTATION

GLU is implemented in R and requires the following R packages: optparse, ggplot2, stringr (see GitHub repository [https://github.com/MRCIEU/GLU] for package versions). A preprocessing step converts device generated CGM data to the CSV format required by GLU (detailed in the GitHub repository; scripts to preprocess Medtronic iPro2 (27) data are included in the GLU repository). Although we have developed GLU using example data from the Medtronic iPro2 device, we have ensured it can be easily used with data from other devices (e.g. Abbott’s Freestyle Libre (28)) by using alternative preprocessing steps. After the initial preprocessing step GLU is run by specifying two directories; the location of the CGM data files, and the location where derived data (e.g. summary variables and plots) should be stored. The CGM data is processed in two main stages: 1) quality control, and 2) deriving summary variables (illustrated in Figure 1). GLU allows the user to specify optional arguments, and these include:

- *nightstart* and *daystart*: Specifies the start time of the day-time and night-time periods of each day to accommodate different populations (e.g. an early bedtime may be more appropriate for studies of children). By default, night-time is between 11.00pm and 6.30am. If other times are used then this should be reported.
- *pregnancy* and *diabetes:* Indicates that the data pertains to pregnant women or diabetic patients, respectively, such that summary variables specific to these populations are derived (i.e. the thresholds used to determine the time spent in hypo-, normo- and hyper-glycaemia levels, described in the ‘Deriving glucose summary variables’ section below). If neither of these options are selected summary variables are produced that assume participants are from a ‘general population’ without selection for pregnancy or diabetes.
- *impute*: Specifies that GLU should perform ‘approximal’ imputation, rather than restricting to ‘complete days’, as described in the ‘CGM data quality control’ section below.

GLU generates a comma-separated value (CSV) file of derived summary variables, which can be imported into statistical software for analysis.

### CGM data quality control

GLU performs quality control to help researchers ensure the integrity of the data, consisting of 3 automated steps: resampling, outlier identification and dealing with missing data (illustrated in Supplementary figure 1). GLU also provides plots for manual review of the CGM data after these automated steps.

#### Resampling

We resample the sensor glucose values across each participant’s CGM sequence to one-minute intervals using linear interpolation (i.e. assuming a straight line between values at adjacent time points), to facilitate computation of summary variables. Given two adjacent time points *t1* and *t2*, with sensor glucose values *SG1* and *SG2*, respectively, linear interpolation estimates the glucose value of time point *t*′ *t*_1_ ≤ *t*′ ≤ *t*_2_ as:

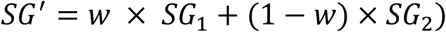

where 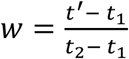.

#### Outlier detection

Previous work has suggested that outliers can be detected by identifying time points that are more than two standard deviations (SD) from the sensor glucose values at both the previous and subsequent time points (5). However, as noted previously (6,19,29), glucose levels may not be normally distributed, so SD may not be an appropriate measure of variability. Furthermore, this approach is sensitive to the resolution of the glucose trace such that changes in resolution would affect which regions of a glucose trace are marked as outliers. This is because SD is invariant to changes in sampling frequency of a glucose trace, while the difference in glucose levels between adjacent time points is not. For example, if sensor glucose is recorded every minute rather than every 5 minutes then the difference in glucose between adjacent time points will be smaller but the overall distribution of sensor glucose values, and hence the outlier detection threshold (based on the SD of this distribution), will not change.

Using data described in our usage example (see Usage section below), we visually assessed the distribution of sensor glucose values for each participant and found these distributions to be very variable – some were normally distributed while others were skewed. We therefore base our outlier detection on the distribution of the differences of adjacent sensor glucose values rather than the distribution of sensor glucose values. We found that the distributions of the difference of adjacent sensor glucose values were more consistently normally distributed compared with the distributions of sensor glucose values. Also, using the differences of adjacent values means that this approach is invariant to changes in the resolution of a glucose trace. We use a threshold *d*, of *k*SD of a participant’s distribution of differences between adjacent values (30). Time points with a glucose value that deviate more than *d* from the value at both the previous and subsequent time points, are marked as outliers for further consideration by the researcher. We chose a threshold of 5SD based on experimentation with our example data (see Supplementary section S1 for further details). Users can also change the value of *k* using GLU’s *outlierthreshold* argument (see GLU GitHub repository for details), to make the outlier detection more conservative or lenient. Should outliers be detected and confirmed by visual inspection of the glucose trace then researchers may wish to: 1) use other data such as diet diaries to determine whether detected outliers may be due to some underlying cause such as food intake (rather than erroneous), and 2) perform sensitivity analyses to see the effect that removing identified outliers has on their results. Our outlier detection method uses a threshold determined using artificial outliers because we have no CGM data containing clear (erroneous) outliers on which to base our approach (Supplementary section S1). As CGM becomes more widely used, it will be possible to improve detection of outliers using outlier examples, and we plan to update GLU outlier detection as the field matures.

#### Assessing the impact of missing data assumptions

CGM data may have missing time periods when the device is unable to record an interstitial glucose value, for example, if the device becomes displaced. When missing periods do exist, there may be systematic differences between the missing and observed values in the CGM data, such that the derived GLU summary variables may be biased. For instance, if sensor displacement (or removal) occurs during swimming and swimming is associated with low glucose values, then a swimmer’s average glucose levels estimated using the observed data may be higher compared to the true underlying value. Under those circumstances associations of the GLU summary variables with a potential outcome or a risk factor may be biased. Alternatively, the CGM missing time periods may be missing completely at random – for instance, some technological failures of CGM devices may be due to chance. We note that there are two related but distinct biases when using GLU derived summary variables: 1) bias of the derived values of participants GLU summary variables, and 2) bias of subsequent analyses using these summary variables. Bias from the former does not necessarily cause bias for the latter as this depends on the specific analyses performed.

GLU provides two approaches to help address missing data called ‘complete days’ and ‘approximal imputed’, that make different missingness assumptions. GLU’s complete days approach uses only days with complete sensor glucose values to derive glucose characteristics (e.g. 24(hours)*60/5=288 values when using CGM data with 5 minute intervals). If the days of CGM data are missing completely at random (MCAR_days_) such that there are no systematic differences between the days with and without missing CGM data, then the derived CGM statistics will be unbiased, hence this missingness will not bias results of subsequent analyses (31). The MCAR_days_ assumption of the complete days’ approach may be violated. For example, characteristics of the participants such as their age or employment status may influence whether or not they complete the required number of capillary blood tests or the likelihood of the CGM device being displaced. However, even when MCAR_days_ does not hold analyses using GLU’s complete days statistics may still be unbiased depending on the specific further analysis in which they are used (31).

In general, imputation may help to reduce the amount of excluded data and relax the missing data assumptions, such that missing at random (MAR) (or sometimes missing not at random [MNAR]) may be assumed rather than MCAR (31). However, glycaemic control is influenced by several characteristics such that imputing portions of a glucose trace is non-trivial. When a participant’s diet and exercise are identical across different days then time-matched data from other days can be used to mean impute missing time points (5), but in most epidemiological studies where data are collected passively under free-living conditions this is not appropriate.

GLU includes a simple imputation approach that fills in the missing periods using nearby data. We refer to this as ‘approximal imputation’. Our approach splits the missing period in half, and uses the sensor glucose data on the left to fill in the left half, and the sensor glucose data on the right to fill in the right half, as illustrated in Supplementary figure 2. Formally, given a missing period of sensor glucose values *{SG_i_, SG_i+1_ … SG_j-1_, SG_j_}, {SG_i_ … SG_k_}* is replaced with *{SG_i-k-1_ … SG_i-1_}* and *{SG_k’_ … SG_j_}* is replaced with *{SG_j+1_ … SG_2j+1-k’_}*, where *k=floor((j-i)/2)* and *k’=k+1*. Each missing sequence must be less than 6 hours long to be considered for imputation. The imputed data is labelled such that the transitions between non-imputed and imputed sections, and transitions between the left and right halves of imputed sections, are not incorporated into summary variables – i.e. only transitions within sections are incorporated (see Supplementary figure 2).

Approximal imputation may help to reduce bias in the derived CGM statistics and hence bias in subsequent analyses that use these statistics. Under the assumption that nearby regions of each missing period are representative of that particular missing period, then CGM statistics derived from approximal imputed data may be less biased. It may however be more likely that missing regions are systematically different to their nearby non-missing regions. For example, if a device is unable to record very high glucose values then nearby glucose values will be systematically lower than the missing region. In this case approximal imputation may still help to reduce bias in the derived CGM statistics. This is because, if days with missing data are systematically different to days without missing data then approximal imputation will enable information from (the non-missing time periods on) these systematically different days to be incorporated into the derived summary variables. Similarly, if the CGM data are MCARday then the summary variables derived using approximal imputed data will be unbiased and more precise than the complete days version.

By default, GLU uses the complete days approach. Users can use the approximal imputation approach by running GLU with the *impute* argument. A researcher wishing to apply another imputation approach to their data (e.g. mean imputation, if appropriate) can do this prior to running GLU. In the rest of this paper we refer to days with complete CGM sequences (after imputation if this option is used) as the set of *included* days. We would suggest that researchers run their analyses using both complete days and approximal imputation and present all results from further analyses (one set could be in supplementary material) so that over time we can learn more about the nature of CGM missing data and its impact on different research questions.

#### Manual review

In the Data visualisation section we describe two plots generated by GLU; these can be used to further check data validity (see Usage section for a description of how we do this in our example).

### Deriving glucose summary variables

After quality control, GLU derives a set of summary variables illustrated in Figure 1. A full list of summary variables computed by GLU is given in Supplementary table 2. Each characteristic is calculated for each day in a participant’s CGM data and, where appropriate, each day-time and night-time period separately (see Supplementary table 2) (8). GLU also provides the average of each summary variable across all days for each participant to give a single overall value for each summary variable (for each participant). For example, GLU returns the following AUC statistics: a) AUC for each included day (24-hours), b) AUC for each included day for the night-time period, c) AUC for each included day for the day-time period, d) mean AUC over all included days (based on a 24-hour day), e) mean AUC of night-time periods over all included days, and f) mean AUC of day-time periods over all included days. The daily statistics provided by GLU allow variability both between and within days to be assessed.

Glucose summary variables output by GLU were chosen to represent broad categories of glucose characteristics that reflect a set of 6 broad domains that might, independently of each other, relate to outcomes or be influenced by exposures (including interventions in RCTs). Supplementary table 1 lists these variables, together with other variables that have been included in some publications but are not included here (together with our reasons for not including them). The 6 broad domains are: overall glucose levels, overall variability (dispersion), excursions (deviations from ‘normal’), variability from one moment to the next, fasting levels and post-event levels. GLU includes one variable from each domain and brief explanations for these choices are given in Supplementary table 1. For example, we considered three measures of dispersion that have been used in previous publications – standard deviation (SD), coefficient of variation (CV) and median absolute deviation (MAD); GLU includes only MAD. This is because sensor glucose values may not be normally distributed and the number of sensor glucose values across which GLU will calculate dispersion will be low (e.g. 1 day contains 288 values for 5 minute epochs), meaning that SD and CV are unlikely to be credible measures of dispersion. All GLU summary variables are independent of the length of the time period for which it is calculated.

#### Overall glucose levels

Overall glucose levels are characterised by the AUC, and specifically GLU derives the mean AUC per minute so that these levels are comparable across time periods of different lengths (e.g. night-time versus day-time) (8). For each day, the AUC is calculated using the trapezoid method (5), as the sum of the area of the trapezoids created using linear interpolation between sensor glucose values at adjacent time points (as described above). We divide by the number of minutes in the time period (e.g. 1440 for whole days) to give the average glucose (mmol/L) per minute.

#### Proportion of time in hypo-, normo- and hyper-glycaemia

We calculate the proportion of time spent in hypo-glycaemia, normo-glycaemia and hyper-glycaemia (8,20,32). In a ‘healthy’ (non-diabetic) and non-pregnant population hypo-glycaemia is defined as <3.3 mmol/L and hyper-glycaemia as ≥10mmol/l (33). The default output from GLU uses these thresholds to define hypo- and hyper-glycaemia in healthy non-pregnant populations (with normo-glycaemia defined as ≥3.3 to <10mmol/L). In patients with diabetes GLUs default for hypo-glycaemia is <3.9 mmol/L and for hyper-glycaemia is ≥10.0 mmol/L (with normo-glycaemia defined as ≥3.9 mmol/L to <10mmol/L) (34). For ‘healthy’ (non-diabetic) pregnant women we use the UK National Institute for Health and Care Excellence (NICE) recommended target range during pregnancy, ≥3.9 mmol/L to <7.8 mmol/L to define normo-glycaemia, and <3.9mmol/L and ≥7.8mmol/L to define hypo- and hyper-glycaemia (35). As already described, these diabetic and pregnancy specific thresholds can be specified using GLU’s *diabetic* and *pregnancy* arguments, respectively. Because thresholds for defining hypo- and hyper-glycaemia (in ‘healthy’, diabetic and pregnant populations) vary geographically and over-time, (35,36), and differ for other groups (for example patients on intensive care units (20)), GLU also allows users to specify other thresholds. However, since GLU is intended to provide standard measures that can be compared (and as appropriate pooled) across studies, where researchers do this a clear justification should be given.

#### Overall variability

While SD and CV are widely used measures of glucose variability (9,25), as discussed above, the distribution of sensor glucose values for a given participant may not be normally distributed. For this reason we use the MAD as a measure of overall variability of sensor glucose levels, defined as:

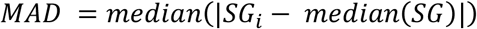

Thus, after calculating the distance of each sensor glucose value from the median value, MAD is the median of these distances.

#### Variability from one moment to the next

We capture variability in a person’s glucose levels across time using a measure based on the length of the line of a glucose trace (i.e. as if the peaks and troughs were stretched out into a line). This idea was recently suggested for CGM data (37) and previously proposed as a measure of complexity for time-series analyses in general (38). Intuitively, if you stretch out a glucose trace then the resultant straight line will tend to be longer when a trace has a larger overall variability (represented by MAD) and is more complex (a higher number of peaks, valleys and values (38)), see Supplementary figure 3 and (38) for examples. This is distinct from MAD because, unlike MAD, the length of the line is affected by the order of the sensor glucose values, i.e. how the sensor glucose values change from one moment to the next (see Supplementary figure 4). Glycaemic variability percentage (GVP) (37) is a rescaling of the average length of the line per minute such that a trace with no variability (i.e. a constant trace) has a GVP of zero. A trace with a GVP of 100% would imply that the length of the trace is double the length of a straight glucose trace. We adapt this measure to capture complexity but not overall variability (in line with (38)), as overall variability is captured by MAD. We standardise each glucose trace prior to deriving GVP by subtracting the median and dividing by the MAD. We refer to the GVP calculated using the standardised glucose levels as standardised GVP (sGVP). Formally, sGVP is defined as:

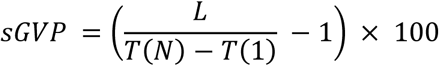

where

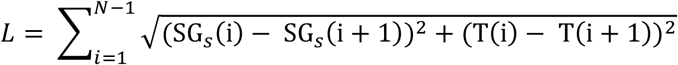

and T and SG_s_ are vectors of timestamps and *standardized* sensor glucose values, respectively, of length *N* (38).

This measure satisfies three useful properties: 1) invariance to the intervals between time points, 2) invariance to differences in overall variability and 3) invariance to differences in the duration of the CGM trace. The first property is satisfied by using a measure based on the length of the line (see Supplementary figure 5a) and means that results of work using different intervals can be compared or meta-analysed. The second property is satisfied by standardising the CGM trace before calculating GVP (see Supplementary figure 5b) and means that associations with sGVP are not due to a relationship with overall variability (i.e. MAD). The third property is satisfied by dividing by the total duration in the above equation (see Supplementary figure 5c) and means that variability from one moment to the next can be compared across time periods of different length.

#### Fasting glucose proxy

While fasting glucose has previously been reported using CGM data, the methods used to derive this measure can be unclear (39,40). In studies where meal times are known fasting glucose levels may be inferred using CGM data recorded before breakfast or after at least 7 hours fasting (5,21,41). For example, using the mean of the 6 consecutive values (with 5 minute intervals) before breakfast (21). Others have used glucose levels during particular periods of the night-time as fasting levels, when meal times are not known (42). This can be problematic if participants eat during the night-time period (5), which occurs in an important minority who may be different in terms of their health and health related behaviours to those who do not eat during the night (43). GLU derives a general proxy measure of fasting glucose that does not require knowledge of meal times, calculated as the mean of the 30 lowest consecutive minutes (equating to 6 CGM values at 5 minute intervals) during the night-time.

#### Event statistics

Studies may ask participants to report their meal times and where this is the case GLU will generate 3 statistics describing subsequent glucose levels: time to peak, and glucose levels 1- and 2-hrs post-prandial (5).

Time to peak is calculated as the number of minutes from the meal to the next sensor glucose peak– i.e. the nearest subsequent sensor glucose value *SG*_*t*_ at time t where *SG*_*t1*_<*SG*_*t*_> *SG*_*t*2_, and *t*1 and *t*2 are the nearest previous and subsequent time points to t, respectively, where *SG*_*t*_ ≠ *SG*_*t*1_ and *SG*_*t*_ ≠ *SG*_*t*2_. We cannot simply find the time point with a higher glucose value than the time points directly before and after, as the peak may consist of a plateau where multiple time points have the same value.

The 1-hr and 2-hr post-prandial glucose measures are calculated as the AUC during the 15-minute period 1- and 2-hrs, respectively, after the meal was recorded. We also calculate the 1- and 2-hr AUC for exercise and medication events, when this information is available.

### Data visualisation

The following plots are produced by GLU:

- Sensor glucose trace plots for all participants that can be visually inspected. This plot also includes indicators of events (where these are provided) including the timing of meal, exercise, use of relevant medications and capillary blood glucose measurement levels. Identified outlier values and imputed time periods (as described above) are also shown on these plots.
- Poincare plots to illustrate the stability of each participants blood glucose levels (10,29,32). Each point on a Poincare plot is the sensor glucose level at time point *t* (on the x-axis) against the sensor glucose level at time point *t* + 1 on the y-axis. Thus, where a participant’s sensor glucose levels change slowly their Poincare plot will be aligned along the ascending diagonal, but those with erratic (and potentially erroneous) sensor glucose levels will have a spread further from the ascending diagonal.

Example sensor glucose trace plots and Poincare plots are shown in Supplementary figure 6, and in the GLU GitHub repository.

## USAGE

In this example, we demonstrate GLU by deriving GLU summary variables from CGM data measured during pregnancy and postnatally, and exploring associations of BMI with these variables during pregnancy.

### Study sample

We used data from the Avon Longitudinal Study of Parents and Children-Generation 2 (ALSPAC-G2). The ALSPAC study website contains details of the data that are available through a fully searchable data dictionary: [http://www.bris.ac.uk/alspac/researchers/data-access/data-dictionary/]. The original ALSPAC cohort (women recruited during an index pregnancy in the early 1990s; ALSPAC-G0) and their index children (ALSPAC-G1) have been described in full elsewhere (1,2). ALSPAC-G2 refers to the children of ALSPAC-G1 and recruitment to this cohort began in June 2012, and further information can be found at http://www.alspac.bris.ac.uk. The data presented here come from a pilot study of CGM in pregnant/postnatal women, which began recruiting ALSPAC-G1 women (or the female partners of ALSPAC-G1 men) during their pregnancy in February 2016. These women were invited to wear a Medtronic iPro2 CGM on their buttock, abdomen or arm, for 5 days, at up to four time points: in early (<28 weeks gestation [median=21 weeks gestation, IQR=(18, 23) range=(6, 27)]) and late (>=28 weeks gestation [median=34 weeks gestation, IQR=(32, 35) range=(28, 36)]) pregnancy, and 6- and 12-months postnatal [median=28 weeks, IQR=(26, 31) and median=58 weeks, IQR=(55, 63.5), respectively]. We refer to the CGM data collected at a particular time point for a particular participant as a CGM instance. While wearing the device, participants were asked to measure their capillary blood glucose by finger prick 4 times daily, for CGM calibration, and record mealtimes in a hand-written diary.

In this pilot a total of 96 CGM instances had been collected, in 63 women. Using GLU’s complete days approach, nine of the 96 instances were excluded due to missing data (one recorded no sensor glucose data and eight had no complete days). One participant has two early pregnancy time points corresponding to two different pregnancies; we excluded the time point for the later pregnancy. We also excluded one participant (with 1 time point) who did not have a measure of BMI. Thus, our pilot includes 85 CGM instances (including a total of 319 included days) – 29 in early pregnancy, 25 in late pregnancy, 15 at 6 months postnatal and 16 at 12 months postnatal. These 85 instances were measured in 61 women. Imputing the sensor glucose data resulted in an additional 2 CGM instances with at least 1 included day, for 2 additional participants. Hence, our imputed dataset includes 87 CGM instances, with a total of 334 included days, in 63 women. Supplementary table 3 shows the patterns of repeat instances within our sample, using both the complete days and approximal imputed approaches.

Women’s weight and height were measured at the clinic visit when the CGM device was inserted and used to calculated body mass index (BMI; kg/m^2^). We considered age, parity and gestational age at CGM measurement as potential confounding factors. Age and parity were reported by the woman; gestational age was calculated from the dates for which the CGM was worn and the woman’s expected date of delivery based on her antenatal records (for the vast majority this would be based on a dating scan).

### Analyses

Since GLU uses different thresholds for defining hypo- normo- and hyper-glycaemia in pregnant compared with non-pregnant women, we divided our CGM instances into pregnancy and postnatal subsets. For the pregnancy subset, we ran GLU with the *pregnancy* argument. For the postnatal subset, we used the default GLU settings (i.e. we didn’t specify any optional parameters). For both, we ran GLU with the complete days approach (which is used by default), and approximal imputation approach (by specifying the *impute* argument). We manually reviewed the trace and Poincare plots to determine whether there may be any anomalies. Poincare plots show how a person’s glucose varies across moments in time (specifically one minute to the next, because GLU resamples CGM data to 1-minute intervals as a pre-processing step). A deviation from the trend along the ascending diagonal on this plot may reflect an erroneous sensor glucose value in the original CGM data, rather than true variation of glucose levels. Sensor glucose values will tend to vary smoothly on CGM trace plots so erratic changes shown on these plots may also indicate erroneous data.

We summarised our derived GLU summary variables at each of the 2 pregnancy and 2 postnatal time points using median and interquartile range (IQR). We then examined the association between early pregnancy BMI (exposure) and GLU CGM derived variables during pregnancy, using the 43 women with a measure during pregnancy. Of these 43 women 32 had just one set of CGM data during pregnancy (18 early- and 14 late-pregnancy) and 11 had data for both early and late pregnancy. For the main analyses we used early pregnancy data for the 11 participants with data at both pregnancy time points. We also undertook a sensitivity analyses in which we instead used late pregnancy measures for these 11 participants. We used linear regression to estimate the association of BMI with the following glucose trace summary variables: overall mean glucose, MAD, sGVP, fasting glucose proxy, post-prandial time-to-peak, and post-prandial 1- and 2-hr AUC. MAD, sGVP and post-prandial time-to-peak were right skewed and hence log transformed to achieve approximately normally distributions of residuals from the regression model. We converted the proportion of time spent in hypo-, normo- and hyper-glycaemia to the number of minutes, by multiplying these by the number of minutes in the defined period (e.g. 1440 in whole days). We then estimated the association of BMI with these outcomes using negative binomial regression. Our analyses were performed using Stata version 14, and code is available at [https://github.com/MRCIEU/GLU-UsageExample/]. Git tag v0.1 corresponds to the version presented here.

### Results

In this pilot we did not identify any outlier time points (Supplementary figure 6 shows some representative trace plots illustrating their smooth nature). Correlations between summary variables are given in Supplementary table 4. The smallest correlation was between MAD and post-prandial time to peak (Pearson’s r=-0.05 [P=0.78]), while the largest was between proportion of time spent in hypo- and normo-glycaemia (Pearson’s r=-0.98 [P<0.01]). While GLU gives the sGVP summary variable, Supplementary table 4 also includes correlations with GVP (that uses the unstandardized glucose trace) for comparison. MAD was positively correlated with GVP (Pearson’s r=0.83 [P<0.01]) and negatively correlated with sGVP (Pearson’s r=-0.58 [P<0.01]). We hypothesised that this is because a glucose trace with a larger overall variability (characterized by MAD) will on average have a lower frequency (intuitively the bigger the deviation the longer it will take to return from this deviation) resulting in a shorter ‘length of the line’ of the glucose trace after standardization. To check this, we derived a simple measure of the number of peaks in each trace and found this had a negative correlation with MAD (Pearson’s r=-0.30 [p=0.05]). Overall, sGVP was less correlated with other GLU summary variables, compared with both MAD and (unstandardized) GVP.

Overall mean glucose levels were very similar (∼5mmol/l) across the four-time points (Supplementary table 5). However, these similar overall glucose levels conceal very different patterns of variation in glucose across the four time points and in the day-versus the night-time. MAD were higher during pregnancy (both early and late) than postnatally, and higher during the day-time compared with the night-time. Fasting glucose was, on average, higher 12 months postnatally compared with early pregnancy. While most time was spent normo-glycaemic both during pregnancy and in the postnatal period, the amount of time spent with levels that fulfilled the criteria for hypo-glycaemia was higher during pregnancy compared with postnatally. In interpreting the proportion of time spent in different glycaemic states across these four time periods it is important to remember that we used the *pregnancy* argument for the early and late pregnancy measures but the default (non-pregnancy, ‘healthy’) option for the postnatal measures. Hence different thresholds were used to define hypo-, normo- and hyper-glycaemic ranges in pregnancy versus postnatally. It is possible that in some women pregnancy related glucose changes might persist postnatally, so we repeated the analyses with the pregnancy function applied to the postnatal time points. The proportion of time spent in normo-glycaemia postnatally was lower when using the *pregnancy* argument, because the pregnancy target normo-glycaemia range is narrower. Results were broadly similar when missing data at some time points were imputed using patterns of data at other times (Figure 2 and Supplementary Table 6).

**Figure 2:**
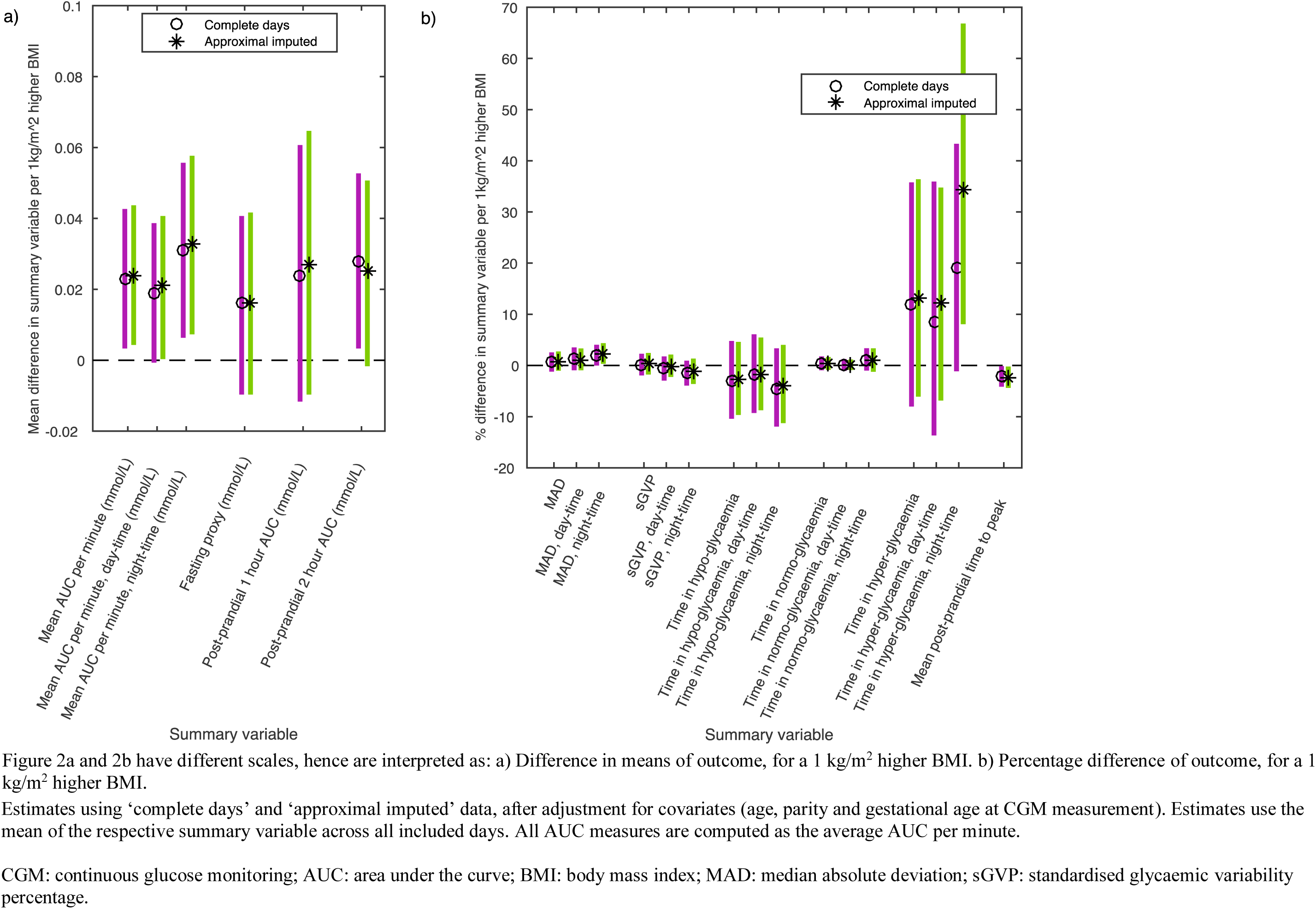
Associations of BMI with GLU summary variables

In analyses using both complete days and approximally imputed data a higher BMI during pregnancy was associated with higher overall mean glucose levels during both the day- and night-time (as measured by AUC), higher time spent in hyper-glycaemia during the night-time, and shorter post-prandial time to peak (Figure 2). For example, during the night-time a 1kg/m^2^ higher BMI was associated with a 0.031 mmol/L higher glucose per minute (95% confidence interval: 0.007, 0.055), after adjusting for covariates (age, parity and gestational age). A higher BMI during pregnancy was associated with higher overall glucose variability during the night-time (as measured by MAD), but we found little evidence of an association with the trace complexity (as measured by sGVP). Results were broadly consistent when we used the late pregnancy measures for the 11 women with both early and late pregnancy results (Supplementary figure 7).

## DISCUSSION

In this paper, we have presented GLU; an open-source tool for researchers working with CGM data. GLU automatically performs quality control, and derives a set of summary variables capturing key characteristics in these data. In contrast to another recently published tool that focusses on analyses specific to the management and detection of diabetes (44), GLU is a general tool for researchers analysing CGM data, and can be applied in ‘healthy’ populations, whether pregnant or not, as well as in studies of people with diabetes. The set of characteristics derived by GLU each represent one of 6 broad domains. For example, overall glucose levels and variability from one moment to the next are represented by the AUC and sGVP, respectively. Our choice of summary variable from each domain was informed by previous work, interpretability of each variable, and the statistical properties of CGM data. For example, given the skewed nature of glucose data GLU uses MAD as a measure of spread, rather than the standard deviation or coefficient of variation (see Supplementary Table 1). GLU also has additional functionality required by researchers, including outlier detection and imputation.

Previously, there has been a lack of consensus in relation to the methods used to derive variables from CGM data. Furthermore, methodological details have often been missing from research articles, making it difficult to replicate studies and compare results across studies (5,23). Using GLU to perform CGM processing and persuading researchers to present findings using all of its summary measures (even if some are in supplementary material) should improve the consistency across studies and hence the opportunity for replication and pooling of results, which is important for improving the robustness of research in this field. Furthermore, over time this would allow insights to emerge related to which glucose trace properties are important for different populations and in relation to different exposures and outcomes. For example, our pilot data suggest that, during pregnancy, BMI is positively associated with mean glucose levels, including during the day and night, and when fasted or postprandial, as well as time spent hyperglycaemic, but has little association with variability or complexity of the glucose trace. We acknowledge that these are pilot data and for some of the outcomes estimates are imprecise (with wide confidence intervals), but it is possible that BMI mostly influences mean levels whereas other exposures might have more influence on glucose variability and complexity. As this field matures we plan to update the tool with any additional summary variables or options that emerge and we encourage researchers to send feedback on the tool and suggest additions (via the corresponding author email).

## ACKNOWLEDGEMENTS

We are extremely grateful to all the families who took part in this study, the midwives for their help in recruiting them, and the whole ALSPAC team, which includes interviewers, computer and laboratory technicians, clerical workers, research scientists, volunteers, managers, receptionists and nurses.

We are grateful to Medtronic Limited who provided the iPro2 continuous glucose monitors and related equipment used in the ALSPAC-G2 pilot study used in this paper (A 1207517).

## AUTHOR CONTRIBUTIONS

LACM conceptualized the tool, developed GLU, contributed to the design of GLU, performed all analyses, wrote the first version of the manuscript, critically reviewed and revised the manuscript and approved the final version of the manuscript as submitted. DAL obtained funds for the CGM pilot study, contributed to decisions regarding the content of GLU and its design and development, developed the pilot analysis plan, critically reviewed and revised the manuscript and approved the final version of the manuscript. NP contributed to the design of GLU, critically reviewed and revised the manuscript and approved the final version of the manuscript as submitted. KT contributed to the design of GLU and decisions regarding the statistical analyses used by GLU, critically reviewed and revised the manuscript and approved the final version as submitted. ML manages the ALSPAC-G2 study, contributed to continuous glucose monitored data collection, critically reviewed the manuscript and approved the final version as submitted. PAF contributed to the design of GLU, critically reviewed and revised the manuscript and approved the final version as submitted.

