## Supplementary Information for "Software application profile: GLU: A tool for analysing continuously measured glucose in epidemiology"

SUPPLEMENTARY MATERIAL

TABLE OF CONTENTS

*SUPPLEMENTARY SECTION S1* *3*
Outlier detection 3
*SUPPLEMENTARY TABLES* *4*
Supplementary table 1: Common CGM summary statistics, grouped into related domains, and reasons for inclusion / exclusion in GLU 5
Supplementary table 2: Summary variables output by GLU 7
Supplementary table 3: Number of participants with each combination of time points 8
Supplementary table 4: Correlations between GLU summary variables 9
Supplementary table 5: Summary statistics across time points 10
Supplementary table 6: Associations of derived GLU summary variables with maternal BMI during pregnancy 12
*SUPPLEMENTARY FIGURES* *14*
Supplementary figure 1: Illustration of GLU preprocessing steps 15
Supplementary figure 2: Illustration of 'approximal' imputation approach 16
Supplementary figure 3: Illustration of length of the glucose trace as a measure of complexity 17
Supplementary figure 5: Illustration of properties satisfied by standardised glycaemic variability percentage (sGVP) 19
Supplementary figure 6: Example plots and associated statistics provided by GLU for two example days 20
Supplementary figure 7: Associations of BMI with GLU summary variables, sensitivity analysis using later pregnancy time point for 5 participants with two pregnancy time points 22
*REFERENCES* *25*

SUPPLEMENTARY SECTION S1

Outlier detection

*Creating a dataset containing artificial outliers:* To determine an appropriate outlier detection method we use our Usage data, which was from the ALSPAC-G2 CGM pilot (see Usage section of the main paper). On visual inspection of these data there were no clear outliers (i.e. points that we deemed could be erroneous). We therefore generated a set of artificial outliers for each day in these data, using the following process (illustrated in Supplementary figure 8). First, we randomly selected a pair of glucose data points in the original CGM data (i.e. before resampling to one minute intervals). Second, we checked to see if the absolute difference in sensor glucose values between these selected points was at least 1mmol/L. If the selected points did not have an absolute difference in sensor glucose of at least 1mmol/L then we resampled again until this was achieved, up to a maximum of 100 attempts. Of the 326 days in our data, this limit of 100 attempts was reached for 2 days, where the CGM data points had low variability such that most/all points were within 1mmol/L of each other, and we excluded these days from our CGM outlier dataset. Finally, we swapped the selected points in each pair and denoted each as an outlier that we aimed to detect. Hence our generated outliers are at least 1mmol/L from the 'true' value in these data, but are also a realistic value for this particular day and participant (since we are swapping two values within a day). After carrying out this process for all days in out ALSPAC-G2 pilot data, our generated outlier dataset consisted of 324 days, each with two artificial outliers.

*Outlier detection approach:* As described in the main paper, we detect outliers using the distribution of the differences between adjacent points. We use a threshold $d$, of $k$ standard deviations (SD) of a participant's distribution of differences between adjacent values. Time points with a glucose value that deviate more than $d$ from the value at both the previous and subsequent time points, are marked as outliers for further consideration by the researcher.

*Choice of outlier detection threshold:* To determine an appropriate value of $k$ we use the ALSPAC-G2 pilot data and the artificial outlier dataset derived from it, as described above. An effective outlier detection method should be able to distinguish between the original ALSPAC-G2 data in which we could not identify outliers by visual inspection, and the outlier dataset in which we have generated artificial outliers. We first assessed a threshold of 4 standard deviations (1) but found this to be too lenient – identifying many points as outliers within our original ALSPAC-G2 data. A threshold of 5 standard deviation was sufficiently stringent – of the 324 days for which artificial outliers were generated, our approach (using the 5SD threshold) identified both outliers on 322 days and 1 outlier only on 2 days. Of the 326 days of real CGM data, our approach (incorrectly) identified time points as outliers on 8 days.

SUPPLEMENTARY TABLES

Supplementary table 1: Common CGM summary statistics, grouped into related domains, and reasons for inclusion / exclusion in GLU

| Glucose characteristic | Included in GLU | Justification for inclusion / exclusion |
|---|---|---|
| **Measures of overall glucose levels** | | |
| Area under the curve (AUC) [average levels across time] (2–4) | Included | Represents overall glucose levels and is widely used. |
| **Measures of overall variability (dispersion)** | | |
| Standard deviation (SD) (2,4,5) | Excluded | Median absolute deviation (MAD) is used as a measure of dispersion that is appropriate as CGM data may not be even approximately normally distributed and the number of values across which GLU will calculate dispersion is low (e.g. for CGM data with 5 minute epochs each day has 288 values) (6–8). |
| Coefficient of variation (CV) (2,4,5) | Excluded | |
| Median absolute deviation (MAD) | Included | |
| **Measures of glycaemic excursions** | | |
| Mean amplitude of glycaemic excursions (MAGE) (2,4,5,9) | Excluded | Proportion of time in low etc. is used as these are interpretable measures of time in hypo- and hyper-glycaemia ranges, and have less ambiguity regarding how to calculate this in an automated way (MAGE was historically calculated manually using visual inspection of the CGM data (9)). |
| Proportion of time spent in low, normal, high glucose levels (4,5) | Included | |
| **Measures of fasting glucose** | | |
| Mean of the 6 lowest consecutive glucose values during the night-time (3) | Included | The lowest mean of the 30 minute period over the night-time (equivalent to the mean of the 6 lowest values when CGM data has 5 minute intervals) has been used as a proxy measure for fasting glucose levels (10). |
| **Measures of variability from one moment to the next** | | |
| Lability index (2,3) | Excluded | Lability index is not normalized by the time period length. MARC, lability index and GVP all capture both overall variability (dispersion) and complexity (a higher number of peaks, valleys and values). GVP is an interpretable measure reflecting the length of the line relative to the minimum length for a given time period (11). The general concept of using the length of the line as a measure of time-series complexity was originally proposed by Batista et al. (12). We calculate the GVP on the standardized CGM traces, and refer to this measure as standardized GVP (sGVP). |
| Standardised glycaemic variability percentage (sGVP) (11) | Included | |
| Mean absolute rate of change (MARC) (5) | Excluded | |
| **Measures of variability across days** | | |
| Absolute means of daily difference (MODD) (2,4) | Excluded | MODD may be most useful where days are matched for diet, physical activity and sleep (and insulin if appropriate) since this measure compares glucose values at the same time across days (13). Hence this is excluded because our focus is on characteristics for free-living CGM data. As GLU provides the derived measures for each day, a researcher can use these to calculate variation across days. |
| **Measures of post-event levels** | | |
| 1-hr and 2-hr post-event AUC (applied to meal and exercise events) (3) | Included | We use the 1hr and 2hr post-event AUC as, unlike the mean of 3 consecutive measures, this is independent of the frequency of time points in the CGM data. |

| | | |
|---|---|---|
| Mean of 3 consecutive measures 1-hr and 2-hr post-event (applied to meal and exercise events) (3) | Excluded | Time to peak is also included in GLU, as a measure of speed of post-prandial response. |
| Post-event time to peak (applied to meal events) (2,3) | Included | |

We select summary variables for inclusion in GLU from 5 broad domains included in this table. Criteria we used included statistical properties of CGM data (e.g normality of sensor glucose distributions), whether a variable is appropriate for free-living CGM data, and interpretability.

Supplementary table 2: Summary variables output by GLU

| | **Day-time** | **Night-time** | **Whole days** | **Average across included days** |
|---|---|---|---|---|
| **AUC (mmol/L)** | YES | YES | YES | YES |
| **MAD (mmol/L)** | YES | YES | YES | YES |
| **sGVP (%)** | YES | YES | YES | YES |
| **Proportion of time in hypo-glycaemia** | YES | YES | YES | YES |
| **Proportion of time in normo-glycaemia** | YES | YES | YES | YES |
| **Proportion of time in hyper-glycaemia** | YES | YES | YES | YES |
| **Fasting proxy (mmol/L)** | *NA* | YES (by definition) | *NA* | YES |
| **Post-prandial time to peak (minutes)** | *NA* | *NA* | YES | YES |
| **Post-prandial 1hr AUC (mmol/L)** | *NA* | *NA* | YES | YES |
| **Post-prandial 2hr AUC (mmol/L)** | *NA* | *NA* | YES | YES |
| **Post-exercise 1hr AUC (mmol/L)** | *NA* | *NA* | YES | YES |
| **Post-exercise 2hr AUC (mmol/L)** | *NA* | *NA* | YES | YES |
| **Post-medication 1hr AUC (mmol/L)** | *NA* | *NA* | YES | YES |
| **Post-medication 2hr AUC (mmol/L)** | *NA* | *NA* | YES | YES |

AUC: area under the curve; hr: hour; MAD: median absolute deviation; sGVP: standardised glycaemic variability percentage.

All AUC measures are computed as the average AUC per minute.

NA: not applicable as these measures are related to events that could occur during the day or night, rather than being based on a continuous section of trace that can be separated into day and night.

Supplementary table 3: Number of participants with each combination of time points

| Number of time points | Time points | | | | Number of participants in ‘complete days’ sample | Number of participants in ‘approximal imputed’ sample |
|---|---|---|---|---|---|---|
| | Early pregnancy | Late pregnancy | 6 months postnatal | 12 months postnatal | | |
| 1 | YES | NO | NO | NO | 12 | 13 |
| | NO | YES | NO | NO | 11 | 11 |
| | NO | NO | YES | NO | 7 | 8 |
| | NO | NO | NO | YES | 9 | 9 |
| 2 | YES | YES | NO | NO | 9 | 9 |
| | YES | NO | YES | NO | 4 | 4 |
| | YES | NO | NO | YES | 2 | 2 |
| | NO | YES | YES | NO | 1 | 1 |
| | NO | YES | NO | YES | 2 | 2 |
| | NO | NO | YES | YES | 2 | 2 |
| 3 | YES | YES | YES | NO | 1 | 1 |
| | YES | YES | NO | YES | 1 | 1 |
| | | | | | 61 | 63 |
| Total number of ‘complete days’ samples at time point | 29 | 25 | 15 | 16 | | |
| Total number of ‘approximal imputed’ samples at time point | 30 | 25 | 16 | 16 | | |

One participant had data from two different pregnancies, and we include only their first pregnancy.

Supplementary table 4: Correlations between GLU summary variables

| | AUC | MAD | GVP | sGVP | Proportion of time in hypo-glycaemia | Proportion of time in normo-glycaemia | Proportion of time in hyper-glycaemia | Fasting proxy | Post-prandial time to peak | Post-prandial 1hr AUC |
|---|---|---|---|---|---|---|---|---|---|---|
| AUC | | | | | | | | | | |
| MAD | 0.03 [0.87] | | | | | | | | | |
| GVP | 0.07 [0.64] | 0.83 [<0.01] | | | | | | | | |
| sGVP | -0.15 [0.32] | -0.58 [<0.01] | -0.24 [0.12] | | | | | | | |
| Proportion of time in hypo-glycaemia | -0.75 [<0.01] | 0.53 [<0.01] | 0.40 [0.01] | -0.17 [0.28] | | | | | | |
| Proportion of time in normo-glycaemia | 0.63 [<0.01] | -0.66 [<0.01] | -0.52 [<0.01] | 0.24 [0.13] | -0.98 [<0.01] | | | | | |
| Proportion of time in hyper-glycaemia | 0.34 [0.03] | 0.77 [<0.01] | 0.68 [<0.01] | -0.37 [0.02] | 0.19 [0.23] | -0.39 [0.01] | | | | |
| Fasting proxy | 0.67 [<0.01] | -0.62 [<0.01] | -0.47 [<0.01] | 0.28 [0.07] | -0.87 [<0.01] | 0.87 [<0.01] | -0.29 [0.06] | | | |
| Post-prandial time to peak | -0.24 [0.19] | -0.05 [0.78] | -0.13 [0.47] | -0.20 [0.26] | 0.07 [0.70] | -0.04 [0.82] | -0.17 [0.36] | -0.06 [0.76] | | |
| Post-prandial 1hr AUC | 0.83 [<0.01] | 0.18 [0.32] | 0.13 [0.48] | -0.34 [0.05] | -0.46 [0.01] | 0.34 [0.05] | 0.43 [0.01] | 0.34 [0.05] | -0.24 [0.20] | |
| Post-prandial 2hr AUC | 0.77 [<0.01] | 0.39 [0.03] | 0.36 [0.04] | -0.36 [0.04] | -0.39 [0.03] | 0.23 [0.19] | 0.59 [<0.01] | 0.19 [0.29] | -0.19 [0.31] | 0.78 [<0.01] |

Pearson’s correlation coefficient [P value]. Using ‘complete days’ approach and pregnancy time points (same dataset used in main analysis with BMI).

GVP is included here to show the effect of standardising the CGM when deriving SGVP.

AUC: area under the curve; SG: sensor glucose; BMI: body mass index; CI: confidence interval; min: minute; MAD: median absolute deviation; GVP: glycaemic variability percentage; sGVP: standardised glycaemic variability percentage.

All AUC measures are computed as the average AUC per minute.

N=43

Supplementary table 5: Summary statistics across time points

| | | **Early pregnancy (≤28 weeks gestational age** | | **Late pregnancy (>28 weeks gestational age)** | | **6 months postnatal** | | **12 months postnatal** | |
|---|---|---|---|---|---|---|---|---|---|
| | **Period of day [1]** | **Complete days** | **Approximal imputed** | **Complete days** | **Approximal imputed** | **Complete days** | **Approximal imputed** | **Complete days** | **Approximal imputed** |
| N | | 29 | 30 | 25 | 25 | 15 | 16 | 16 | 16 |
| AUC [1] (mmol/L) | Whole | 4.79 (4.44,5.25) | 4.78 (4.44, 5.25) | 4.96 (4.69, 5.32) | 4.96 (4.70, 5.32) | 5.02 (4.59, 5.27) | 5.03 (4.64, 5.23) | 5.11 (4.92, 5.30) | 5.15 (4.92, 5.30) |
| | Day | 4.92 (4.62, 5.20) | 4.91 (4.63, 5.20) | 5.14 (4.84, 5.47) | 5.14 (4.84, 5.50) | 5.08 (4.64, 5.34) | 5.09 (4.69, 5.27) | 5.23 (5.00, 5.45) | 5.24 (5.00, 5.45) |
| | Night | 4.46 (4.06, 5.00) | 4.50 (4.09, 5.00) | 4.82 (4.37, 5.02) | 4.82 (4.37, 5.02) | 4.79 (4.39, 5.23) | 4.83 (4.44, 5.31) | 4.78 (4.66, 5.22) | 4.78 (4.67, 5.27) |
| MAD (mmol/L) | Whole | 0.50 (0.37, 0.54) | 0.49 (0.37, 0.54) | 0.54 (0.46, 0.62) | 0.54 (0.46, 0.62) | 0.37 (0.27, 0.40) | 0.37 (0.29, 0.40) | 0.35 (0.32, 0.52) | 0.35 (0.32, 0.52) |
| | Day | 0.43 (0.34, 0.51) | 0.44 (0.34, 0.51) | 0.55 (0.42, 0.70) | 0.55 (0.42, 0.70) | 0.30 (0.22, 0.39) | 0.32 (0.23, 0.43) | 0.32 (0.29, 0.42) | 0.32 (0.29, 0.42) |
| | Night | 0.25 (0.20, 0.28) | 0.24 (0.20, 0.28) | 0.29 (0.22, 0.34) | 0.31 (0.22, 0.40) | 0.24 (0.15, 0.35) | 0.25 (0.18, 0.36) | 0.28 (0.20, 0.36) | 0.28 (0.20, 0.36) |
| sGVP (%) | Whole | 0.13 (0.09, 0.19) | 0.13 (0.09, 0.19) | 0.12 (0.09, 0.14) | 0.12 (0.09, 0.14) | 0.24 (0.16, 0.37) | 0.21 (0.14, 0.34) | 0.21 (0.13, 0.28) | 0.21 (0.13, 0.28) |
| | Day | 0.16 (0.11, 0.22) | 0.16 (0.11, 0.22) | 0.12 (0.11, 0.16) | 0.12 (0.11, 0.16) | 0.28 (0.20, 0.35) | 0.25 (0.20, 0.31) | 0.26 (0.17, 0.32) | 0.26 (0.17, 0.32) |
| | Night | 0.35 (0.26, 0.45) | 0.35 (0.22, 0.46) | 0.24 (0.18, 0.32) | 0.24 (0.18, 0.32) | 0.31 (0.20, 0.50) | 0.31 (0.18, 0.45) | 0.31 (0.19, 0.43) | 0.31 (0.19, 0.43) |
| Proportion of time in hypo-glycaemia per day | Whole | 0.11 (0.04, 0.25) | 0.10 (0.04, 0.24) | 0.07 (0.01, 0.18) | 0.07 (0.01, 0.18) | 0.00 (0.00, 0.02)<br>***0.03 (0.00, 0.08)*** [#] | 0.01 (0.00, 0.03)<br>***0.05 (0.01, 0.08)*** [#] | 0.00 (0.00, 0.02)<br>***0.02 (0.00, 0.07)*** [#] | 0.00 (0.00, 0.02)<br>***0.02 (0.00, 0.07)*** [#] |
| | Day | 0.07 (0.02, 0.18) | 0.07 (0.02, 0.18) | 0.06 (0.01, 0.17) | 0.06 (0.01, 0.17) | 0.00 (0.00,0.00)<br>***0.02 (0.00, 0.06)*** [#] | 0.00 (0.00, 0.02)<br>***0.03 (0.00, 0.07)*** [#] | 0.00 (0.00, 0.00)<br>***0.00 (0.00, 0.03)*** [#] | 0.00 (0.00, 0.00)<br>***0.00 (0.00, 0.03)*** [#] |
| | Night | 0.20 (0.05, 0.41) | 0.18 (0.05, 0.40) | 0.10 (0.02, 0.16) | 0.10 (0.02, 0.16) | 0.00 (0.00,0.05)<br>***0.08 (0.00, 0.23)*** [#] | 0.00 (0.00, 0.05)<br>***0.05 (0.00, 0.18)*** [#] | 0.00 (0.00, 0.05)<br>***0.06 (0.00, 0.18)*** [#] | 0.00 (0.00, 0.05)<br>***0.06 (0.00, 0.17)*** [#] |
| Proportion of time in normo-glycaemia per day | Whole | 0.84 (0.75, 0.96) | 0.84 (0.75, 0.96) | 0.91 (0.82, 0.97) | 0.91 (0.82, 0.97) | 1.00 (0.98, 1.00)<br>***0.96 (0.91, 1.00)*** [#] | 0.99 (0.97, 1.00)<br>***0.95 (0.90, 0.98)*** [#] | 1.00 (0.98, 1.00)<br>***0.98 (0.93, 0.99)*** [#] | 1.00 (0.98, 1.00)<br>***0.98 (0.93, 0.99)*** [#] |
| | Day | 0.86 (0.82, 0.97) | 0.90 (0.82, 0.98) | 0.92 (0.83, 0.98) | 0.92 (0.83, 0.98) | 1.00 (1.00,1.00)<br>***0.98 (0.92, 1.00)*** [#] | 1.00 (0.98, 1.00)<br>***0.96 (0.91, 1.00)*** [#] | 1.00 (1.00, 1.00)<br>***0.99 (0.96, 1.00)*** [#] | 1.00 (1.00, 1.00)<br>***0.99 (0.96, 1.00)*** [#] |
| | Night | 0.80 (0.59, 0.94) | 0.82 (0.60, 0.94) | 0.90 (0.84, 0.97) | 0.90 (0.84, 0.97) | 1.00 (0.95,1.00)<br>***0.92 (0.77, 1.00)*** [#] | 1.00 (0.95, 1.00)<br>***0.95 (0.82, 1.00)*** [#] | 1.00 (0.95, 1.00<br>***0.94 (0.82, 0.99)*** [#] | 1.00 (0.95, 1.00)<br>***0.94 (0.83, 0.99)*** [#] |
| Proportion of time in hyper-glycaemia per day | Whole | 0.00 (0.00, 0.00) | 0.00 (0.00, 0.00) | 0.00 (0.00, 0.01) | 0.00 (0.00, 0.01) | 0.00 (0.00, 0.00)<br>***0.00 (0.00, 0.01)*** [#] | 0.00 (0.00, 0.00)<br>***0.00 (0.00, 0.01)*** [#] | 0.00 (0.00, 0.00)<br>***0.00 (0.00, 0.01)*** [#] | 0.00 (0.00, 0.00)<br>***0.00 (0.00, 0.01)*** [#] |
| | Day | 0.00 (0.00, 0.00) | 0.00 (0.00, 0.00) | 0.00 (0.00, 0.02) | 0.00 (0.00, 0.02) | 0.00 (0.00, 0.00)<br>***0.00 (0.00, 0.01)*** [#] | 0.00 (0.00, 0.00)<br>***0.00 (0.00, 0.01)*** [#] | 0.00 (0.00, 0.00)<br>***0.00 (0.00, 0.00)*** [#] | 0.00 (0.00, 0.00)<br>***0.00 (0.00, 0.00)*** [#] |
| | Night | 0.00 (0.00, 0.00) | 0.00 (0.00, 0.00) | 0.00 (0.00, 0.00) | 0.00 (0.00, 0.00) | 0.00 (0.00, 0.00)<br>***0.00 (0.00, 0.00)*** [#] | 0.00 (0.00, 0.00)<br>***0.00 (0.00, 0.00)*** [#] | 0.00 (0.00, 0.00)<br>***0.00 (0.00, 0.00)*** [#] | 0.00 (0.00, 0.00)<br>***0.00 (0.00, 0.00)*** [#] |
| Fasting glucose proxy (mmol/L) | NA | 3.69 (3.42, 4.11) | 3.69 (3.44, 4.06) | 3.93 (3.67, 4.31) | 3.93 (3.67, 4.31) | 4.09 (3.79, 4.75) | 4.20 (3.87, 4.67) | 4.23 (3.91, 4.44) | 4.26 (3.91, 4.48) |
| Post-prandial time to peak (mins) | Whole | 55.25 (42.29, 84.42) [a] | 55.25 (42.29, 84.42) [a] | 61.30 (47.79, 80.75) [b] | 61.27 (47.17, 73.00) [a] | 59.57 (42.50, 69.00) | 54.48 (42.50, 69.00) [c] | 60.19 (36.93, 103.26) | 60.19 (36.93, 103.26) |
| Post-prandial 1hr AUC (mmol/L) | Whole | 5.02 (4.76, 5.45) [a] | 5.02 (4.76, 5.45) [a] | 5.40 (5.09, 5.67) [a] | 5.42 (5.10, 5.84) [a] | 5.19 (4.93, 5.42) | 5.19 (4.93, 5.37) [c] | 5.39 (4.99, 5.71) | 5.39 (4.99, 5.71) |
| Post-prandial 2hr AUC (mmol/L) | Whole | 5.26 (4.78, 5.39) [a] | 5.26 (4.85, 5.39) [a] | 5.35 (5.13, 5.60) [a] | 5.34 (5.01, 5.60) [a] | 5.14 (4.75, 5.32) | 5.14 (4.75, 5.32) [c] | 5.27 (4.98, 5.56) | 5.27 (4.98, 5.51) |

Summary statistics for all GLU summary variables are the median (inter-quartile range).

AUC: area under the curve; SG: sensor glucose; BMI: body mass index; CI: confidence interval; MAD: median absolute deviation; sGVP: standardised glycaemic variability percentage.

'Complete days' summary statistics are based on 85 CGM data samples from 61 women, with 39 women contributing to just one time point (distributed similarly across the four time points) and 22 women with two or more repeat measures. 'Approximal imputed' summary statistics are based on 87 CGM data samples from 63 women, with 41 women contributing to just one time point (distributed similarly across the four time points) and 22 women with two or more repeat measures. See Supplementary table 3 for details of repeat measures.

[#] ***In bold italics and blue:*** Applying pregnancy thresholds instead of non-pregnancy thresholds, for comparison.

Meal event measures could not be calculated for some participants (e.g. because they have no recorded meals on included days or no peak after a recorded meal event) such that these summaries are based on a subset of our sample.
[a] N=21
[b] N=20
[c] N=15

All AUC measures are computed as the average AUC per minute.

Supplementary table 6: Associations of derived GLU summary variables with maternal BMI during pregnancy

| | | Estimates for a 1 $kg/m^2$ higher BMI [95% CI] [4] | | | |
|---|---|---|---|---|---|
| | | 'Complete days' analysis | | Approximal imputed analysis | |
| | **Time of day included** | **Unadjusted** | **Adjusted [5]** | **Unadjusted** | **Adjusted [5]** |
| N participants | | 43 | 43 | 44 | 44 |
| AUC (mmol/L) [1] | Whole day | 0.025 [0.005, 0.045] | 0.023 [0.004, 0.042] | 0.026 [0.006, 0.045] | 0.024 [0.005, 0.043] |
| | Day-time | 0.021 [0.001, 0.040] | 0.019 [-0.000, 0.038] | 0.022 [0.002, 0.041] | 0.021 [0.001, 0.040] |
| | Night-time | 0.035 [0.011, 0.058] | 0.031 [0.007, 0.055] | 0.035 [0.011, 0.059] | 0.033 [0.008, 0.057] |
| MAD (% difference) [3] | Whole day | 0.593 [-0.808, 2.013] | 0.676 [-0.726, 2.098] | 0.767 [-0.645, 2.198] | 0.831 [-0.570, 2.252] |
| | Day-time | 1.191 [-0.648, 3.065] | 1.283 [-0.464, 3.061] | 1.155 [-0.608, 2.950] | 1.175 [-0.475, 2.853] |
| | Night-time | 2.154 [0.490, 3.845] | 2.001 [0.459, 3.568] | 2.473 [0.805, 4.168] | 2.322 [0.777, 3.890] |
| sGVP (% difference) [3] | Whole day | 0.141 [-1.471, 1.779] | 0.147 [-1.487, 1.809] | 0.281 [-1.311, 1.898] | 0.332 [-1.294, 1.985] |
| | Day-time | -0.608 [-2.428, 1.246] | -0.620 [-2.494, 1.291] | -0.207 [-1.890, 1.504] | -0.085 [-1.789, 1.647] |
| | Night-time | -1.516 [-3.509, 0.519] | -1.530 [-3.473, 0.453] | -1.269 [-3.342, 0.849] | -1.154 [-3.147, 0.880] |
| Time in hypo-glycaemia (% difference) [1,2] | Whole day | -3.825 [-10.436, 3.273] | -3.063 [-9.923, 4.318] | -3.622 [-9.876, 3.066] | -2.776 [-9.219, 4.125] |
| | Day-time | -2.701 [-9.292, 4.369] | -1.863 [-8.809, 5.612] | -2.697 [-8.819, 3.835] | -1.865 [-8.281, 5.000] |
| | Night-time | -5.321 [-12.081, 1.958] | -4.575 [-11.483, 2.873] | -4.844 [-11.658, 2.497] | -3.894 [-10.799, 3.546] |
| Time in normo-glycaemia (% difference) [1,2] | Whole day | 0.457 [-0.449, 1.371] | 0.408 [-0.478, 1.302] | 0.412 [-0.416, 1.247] | 0.386 [-0.412, 1.190] |
| | Day-time | 0.158 [-0.556, 0.878] | 0.106 [-0.598, 0.814] | 0.159 [-0.424, 0.746] | 0.128 [-0.435, 0.695] |
| | Night-time | 1.188 [-0.550, 2.957] | 1.167 [-0.537, 2.900] | 1.030 [-0.806, 2.900] | 1.040 [-0.755, 2.868] |
| Time in hyper-glycaemia (% difference) [1,2] | Whole day | 12.230 [-4.655, 32.104] | 11.851 [-7.547, 35.319] | 12.477 [-4.444, 32.395] | 13.263 [-5.620, 35.925] |
| | Day-time | 11.453 [-6.180, 32.400] | 8.430 [-13.212, 35.469] | 11.368 [-5.539, 31.301] | 12.126 [-6.389, 34.302] |
| | Night-time | 35.524 [2.645, 78.936] | 19.125 [-0.659, 42.849] | 38.632 [9.738, 75.133] | 34.370 [8.546, 66.337] |
| Fasting proxy (mmol/L) | NA | 0.017 [-0.007, 0.041] | 0.016 [-0.009, 0.040] | 0.016 [-0.008, 0.041] | 0.016 [-0.009, 0.041] |
| Post-prandial time to peak (% change) [3,7] | Whole day | -1.921 [-3.275, -0.548] | -2.154 [-3.688, -0.597] | -2.020 [-3.421, -0.599] | -2.311 [-3.911, -0.684] |
| Post-prandial 1hr AUC (mmol/L) [6] | Whole day | 0.026 [-0.008, 0.060] | 0.024 [-0.011, 0.060] | 0.029 [-0.006, 0.064] | 0.027 [-0.009, 0.064] |
| Post-prandial 2hr AUC (mmol/L) [6] | Whole day | 0.027 [0.003, 0.052] | 0.028 [0.004, 0.052] | 0.025 [-0.001, 0.050] | 0.025 [-0.001, 0.050] |

Estimates using 'complete days' and 'approximal imputed' data, after adjustment for covariates (age, parity and gestational age at CGM measurement). Estimates use the mean of the respective summary variable across all included days.

AUC: area under the curve; SG: sensor glucose; BMI: body mass index; CI: confidence interval; min: minute; MAD: median absolute deviation; sGVP: standardised glycaemic variability percentage.

[1] Difference in means of outcome, for a 1 $kg/m^2$ higher BMI.

[2] Negative binomial regression.

[3] Log transformed outcome. Estimates are the percentage change of outcome for a 1 $kg/m^2$ higher BMI.

[4] Regression analyses include pregnancy time points only.
[5] Adjusted for gestational age, participant age and parity.
[6] Sample sizes are lower because some participants have no meals: N ‘complete days’ unadjusted = 33, N ‘complete days’ adjusted = 33; N ‘imputed’ unadjusted = 33, N ‘imputed’ adjusted = 33.
[7] Sample sizes are lower because some participants have no meals: N ‘complete days’ unadjusted = 32, N ‘complete days’ adjusted = 32; N ‘imputed’ unadjusted = 33, N ‘imputed’ adjusted = 33.

All AUC measures are computed as the average AUC per minute.

Associations correspond to those presented in Figure 2 in the main paper.

SUPPLEMENTARY FIGURES

Supplementary figure 1: Illustration of GLU preprocessing steps

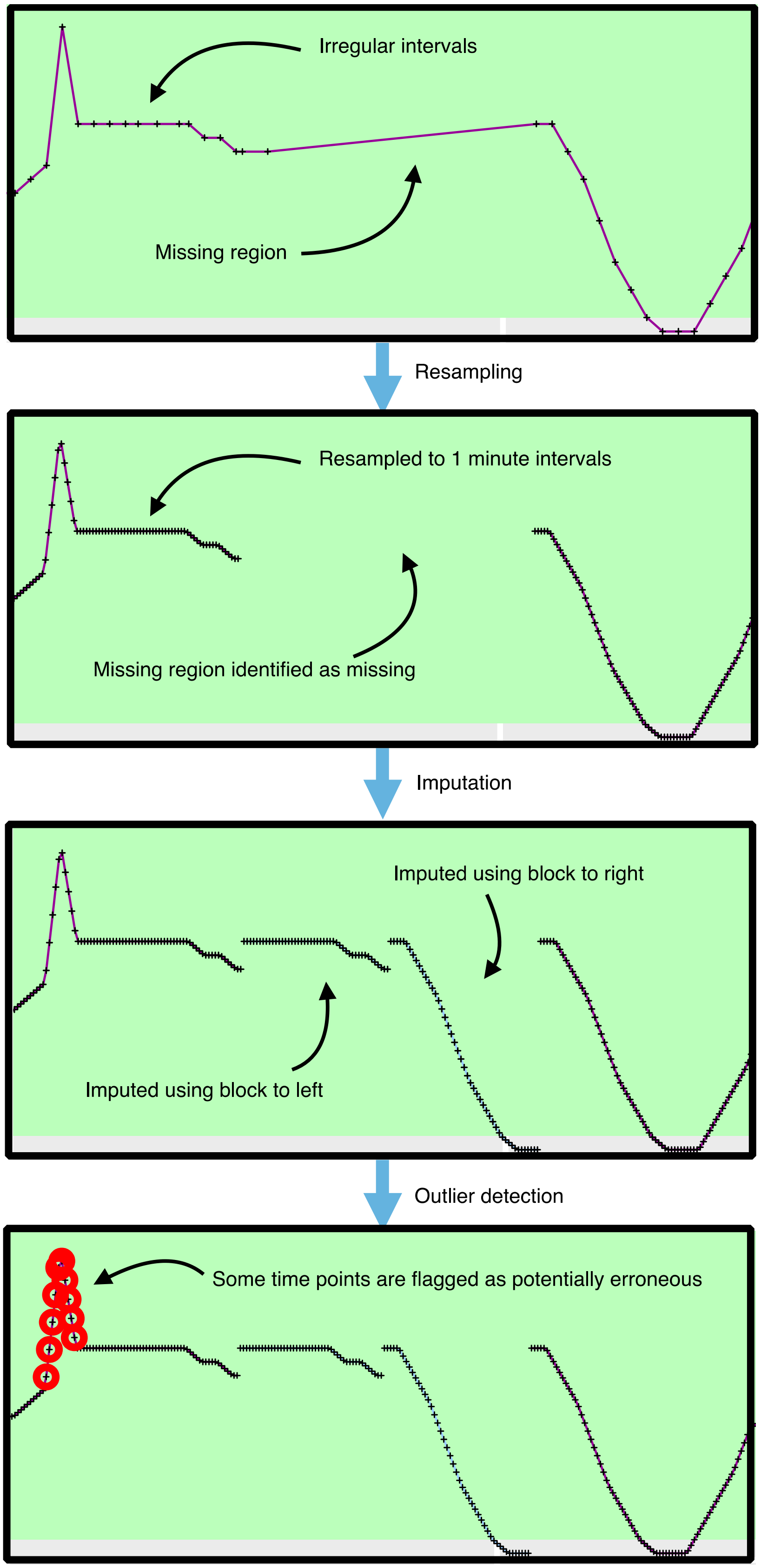

Supplementary figure 2: Illustration of ‘approximal’ imputation approach

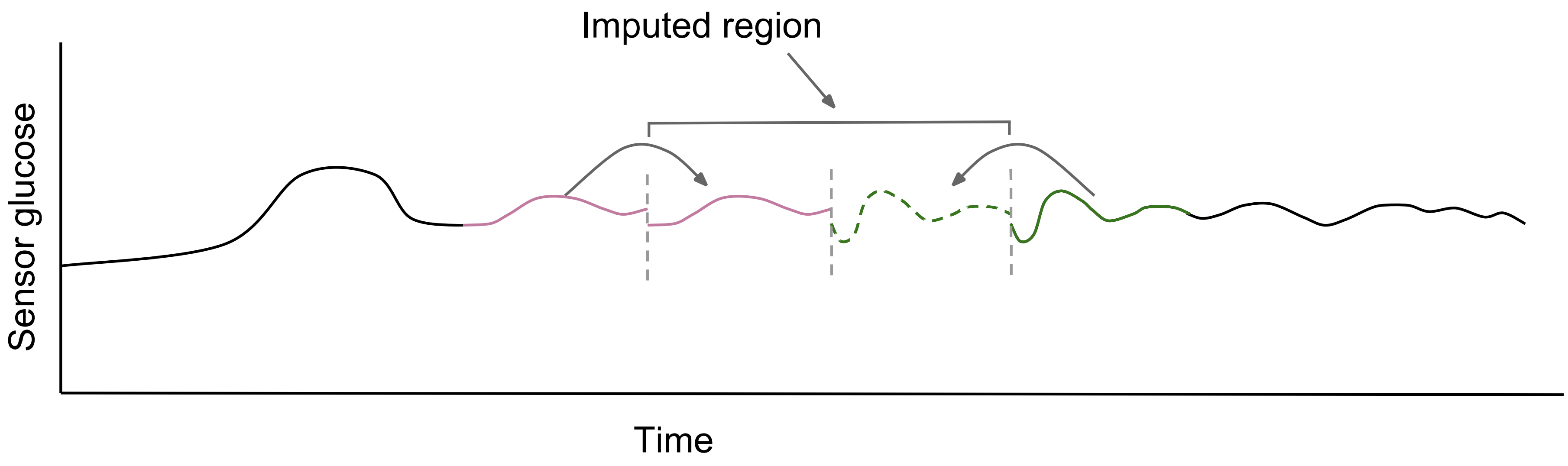


We restrict imputation to missing windows that are at most 6 hours long.

Given a missing window less than 6 hours long, this is imputed using nearby sequences. The left half of the missing window is imputed using the non-missing data to the left of the missing window. The right half of the missing window is imputed using the non-missing data to the right of the missing window.

Dashed vertical lines mark transitions between imputed and non-imputed regions and the left and right parts of an imputed region, and these transitions are not incorporated into GLU summary variables.

Supplementary figure 3: Illustration of length of the glucose trace as a measure of complexity

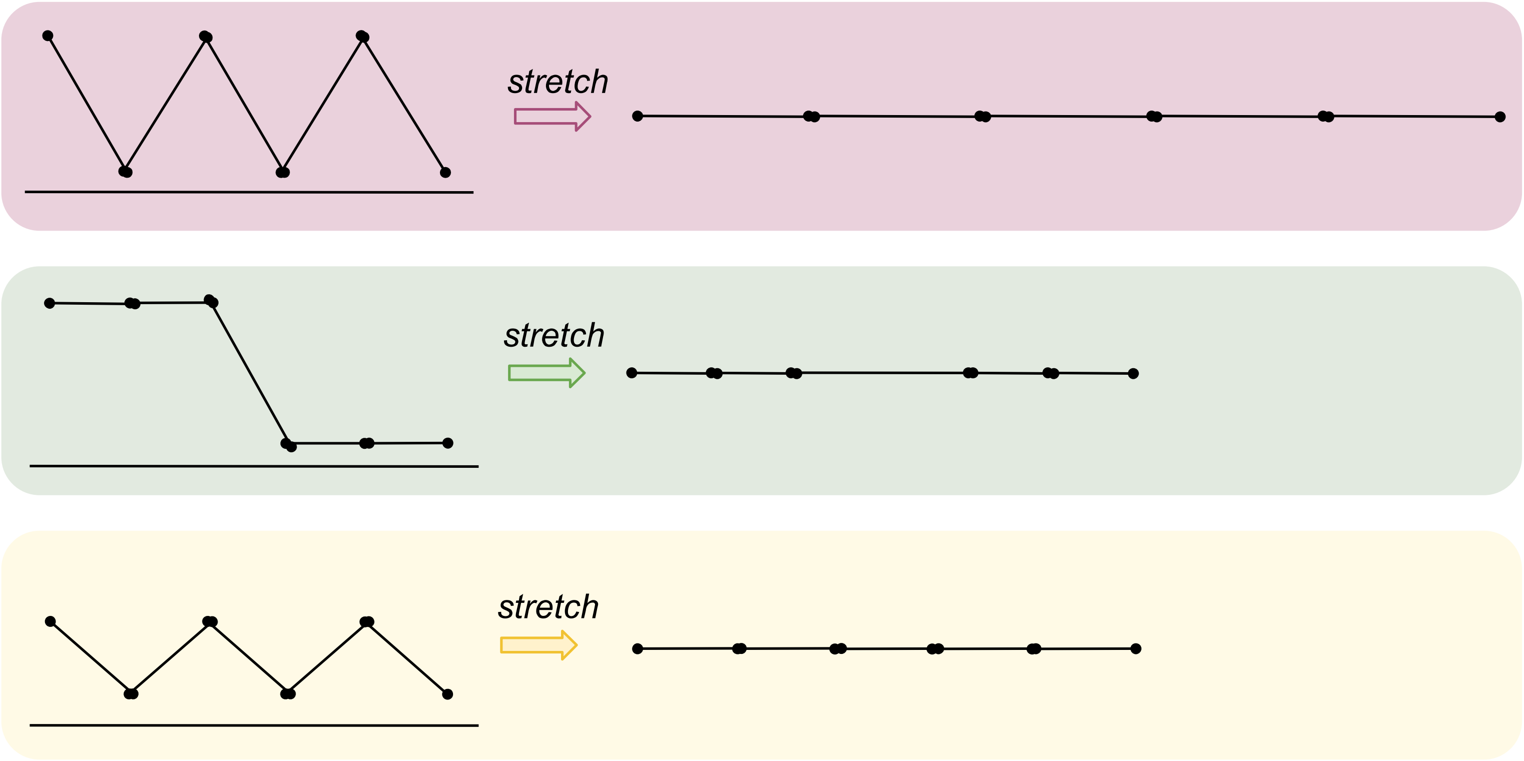


Trace A and B have the same AUC, but trace A has a higher oscillation frequency such that the length of the trace (after stretching) is longer. Trace A has a greater amplitude compared with trace C, and hence the length of the trace (after stretching) is longer. Traces B and C have similar lengths of the trace (after stretching), but different causes; B has a higher amplitude, whereas B has a higher oscillation frequency (trace A instead has constant sections).

Supplementary figure 4: Illustration of the distinction between MAD (representing overall variability) and sGVP (representing variability across moments in time)

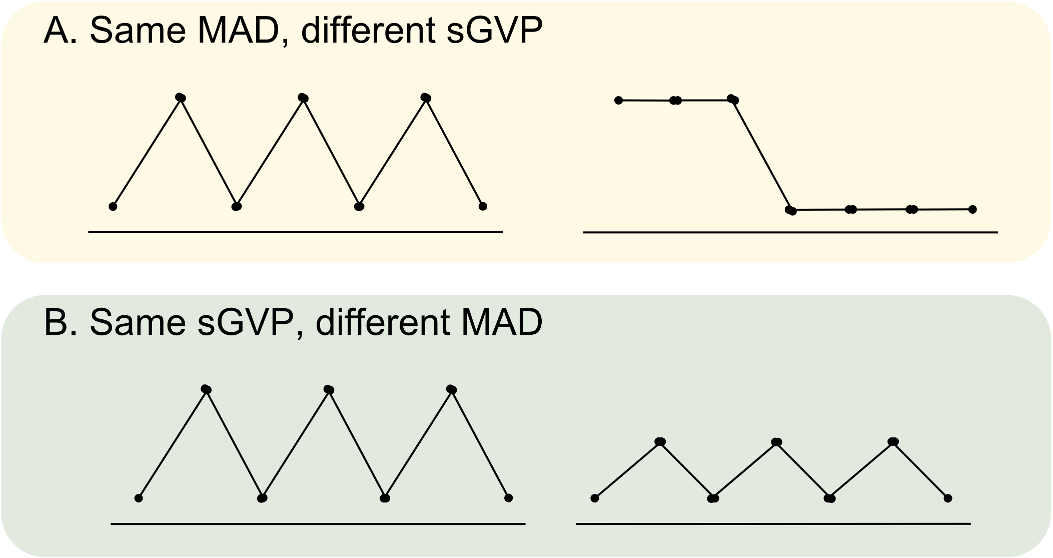


A) Sensor glucose traces have the same MAD (overall variability), with 3 lower and 3 higher values, but different complexity. This demonstrates how the order of the sensor glucose values affects sGVP, but not MAD.
B) Sensor glucose traces have the same sGVP due to the standardisation of sensor values prior to calculating GVP, but different MAD reflecting the difference in overall variability of the two (unstandardized) traces.

Supplementary figure 5: Illustration of properties satisfied by standardised glycaemic variability percentage (sGVP)

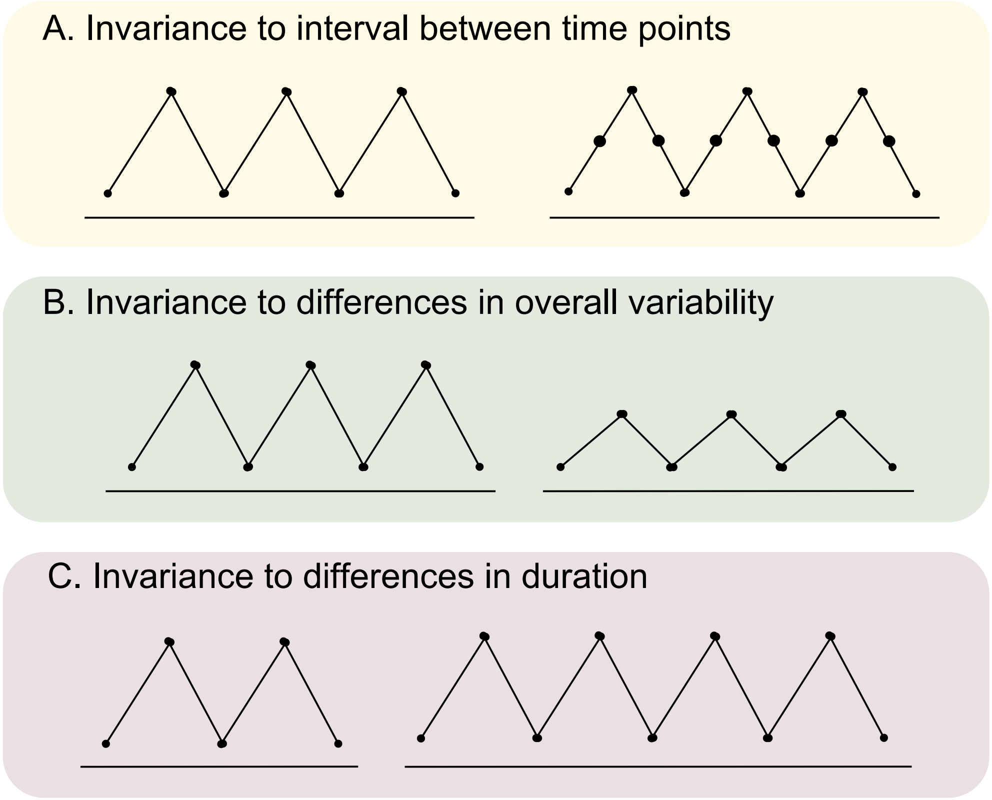


A. The left trace is sampled as half the frequency of the right trace. Two identical traces except for sampling frequency have the same sGVP.
B. These traces are a simple rescaling (of glucose values) to each other. Two traces that are a rescaling of each other have the same sGVP, because the standardised versions will be identical.
C. The time period of the left trace is half the time period of the right trace. As sGVP calculates the length of the line *relative to the shortest line* these traces have the same sGVP value.

Supplementary figure 6: Example plots and associated statistics provided by GLU for two example days

**CGM trace A**

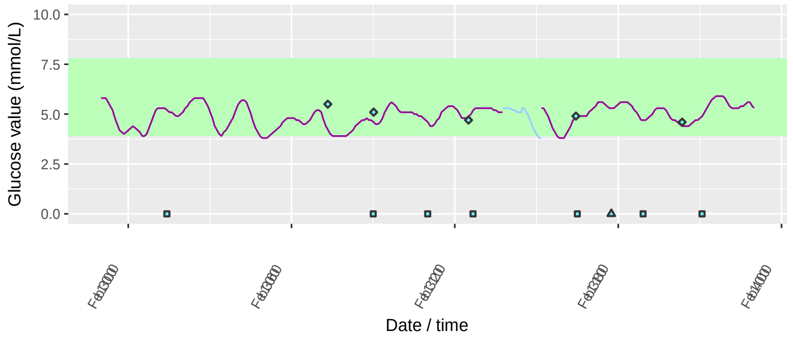


**Poincare plot A**

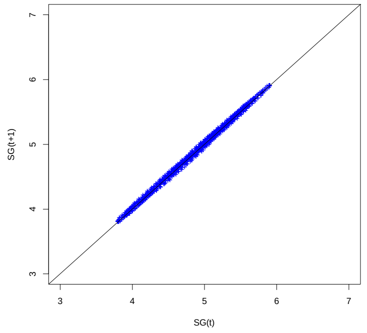


**CGM trace B**

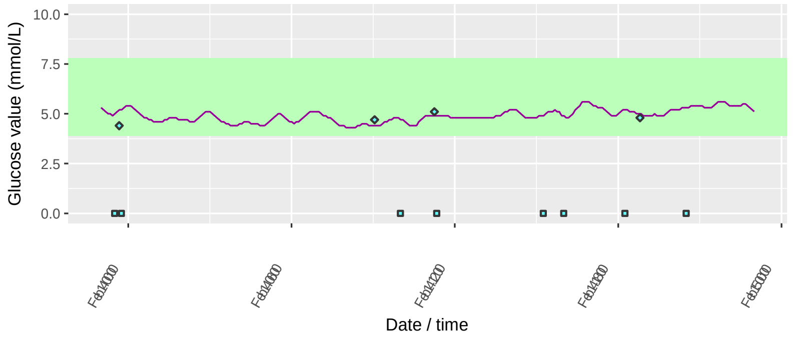


**Poincare plot B**

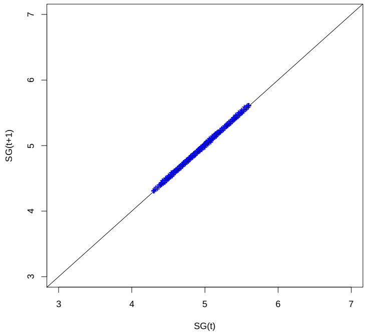


**GLU derived summary variables**

| Summary variable | Day A | Day B |
|---|---|---|
| AUC | 4.90 | 4.91 |
| MAD | 0.35 | 0.21 |
| sGVP | 0.33 | 0.21 |
| Proportion of time in hypo-glycaemia | 0.03 | 0.00 |
| Proportion of time in normo-glycemia | 0.97 | 1.00 |
| Proportion of time in hyper-glycaemia | 0.00 | 0.00 |
| Fasting glucose proxy | 3.87 | 4.43 |
| Post-prandial time to peak (mins) | 37.4 | 59.75 |
| 1 hour post-prandial AUC (avg per minute) | 5.21 | 5.01 |
| 2 hour post-prandial AUC (avg per minute) | 4.88 | 5.04 |
| 1 hour post-exercise AUC (avg per minute) | 4.74 | NA |
| 2 hour post-exercise AUC (avg per minute) | 4.94 | NA |

Event symbols: Diamonds: finger prick blood glucose measurements; stars: medication; triangles: exercise; squares: meals. Default settings are used: night-time start: 11.00; day-time start: 6.30.

AUC: area under the curve; min: minute; MAD: median absolute deviation; sGVP: standardised glycaemic variability percentage.

All AUC measures are computed as the average AUC per minute.

Purple trace line: Sensor glucose values (after resampling to one minute intervals). Blue trace line (in trace A): Imputed section where sensor glucose values were missing in this time period.

Days A and B have similar overall glucose levels as characterised by the AUC.
Day A has higher variability (both overall and across moments in time) than day A as characterised by MAD and sGVP, and the ‘longer’ mass of points on the Poincare plot.
Glucose levels of day B fall completely within the bounds of normo-glycaemia, whereas day A has some time in hypo-glycaemia.
Fasting proxy is lower in day A reflecting the dip in glucose levels around 5AM.
Day B has a longer post-prandial time to peak compared with day A.
Reflecting the steep ascents/descents in day A compared with day B, day A has a higher 1-hr post-prandial AUC but lower 2-hr post-prandial AUC compared with day B.
Day B has no exercise events.

Supplementary figure 7: Associations of BMI with GLU summary variables, sensitivity analysis using later pregnancy time point for 11 participants with two pregnancy time points

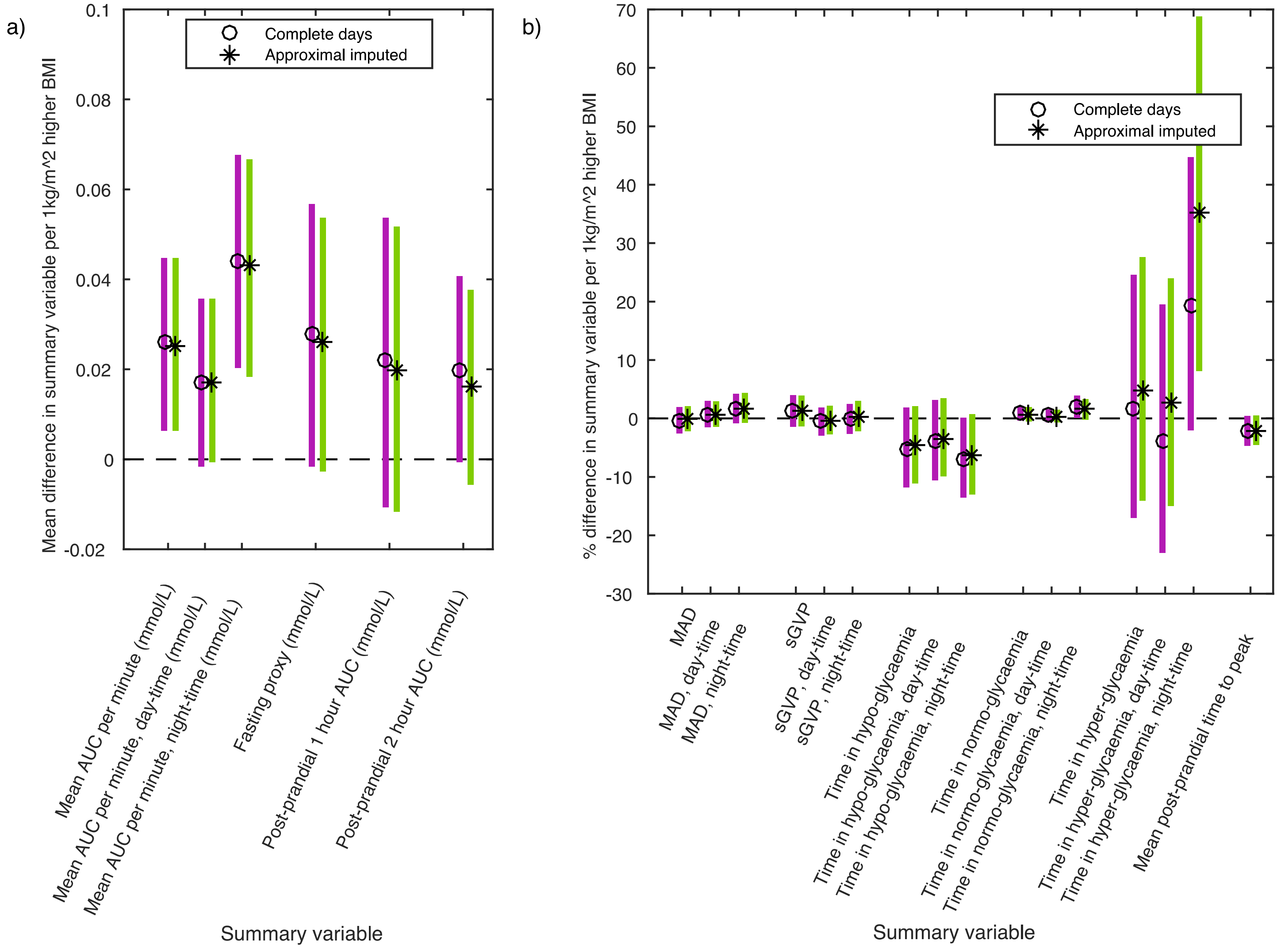

Estimates using ‘complete days’ and ‘approximally imputed’ data, after adjustment for covariates (age, parity and gestational age at CGM measurement). Estimates use the mean of the respective summary variable across all included days.

Analyses are adjusted for age, parity and gestational age.

AUC: area under the curve; BMI: body mass index; MAD: median absolute deviation; sGVP: standardised glycaemic variability percentage.

A) Difference in means of outcome, for a 1 $kg/m^2$ higher BMI.
B) Percentage difference of outcome, for a 1 $kg/m^2$ higher BMI.

All AUC measures are computed as the average AUC per minute.

Supplementary figure 8: Illustration of our approach to generating artificial outliers

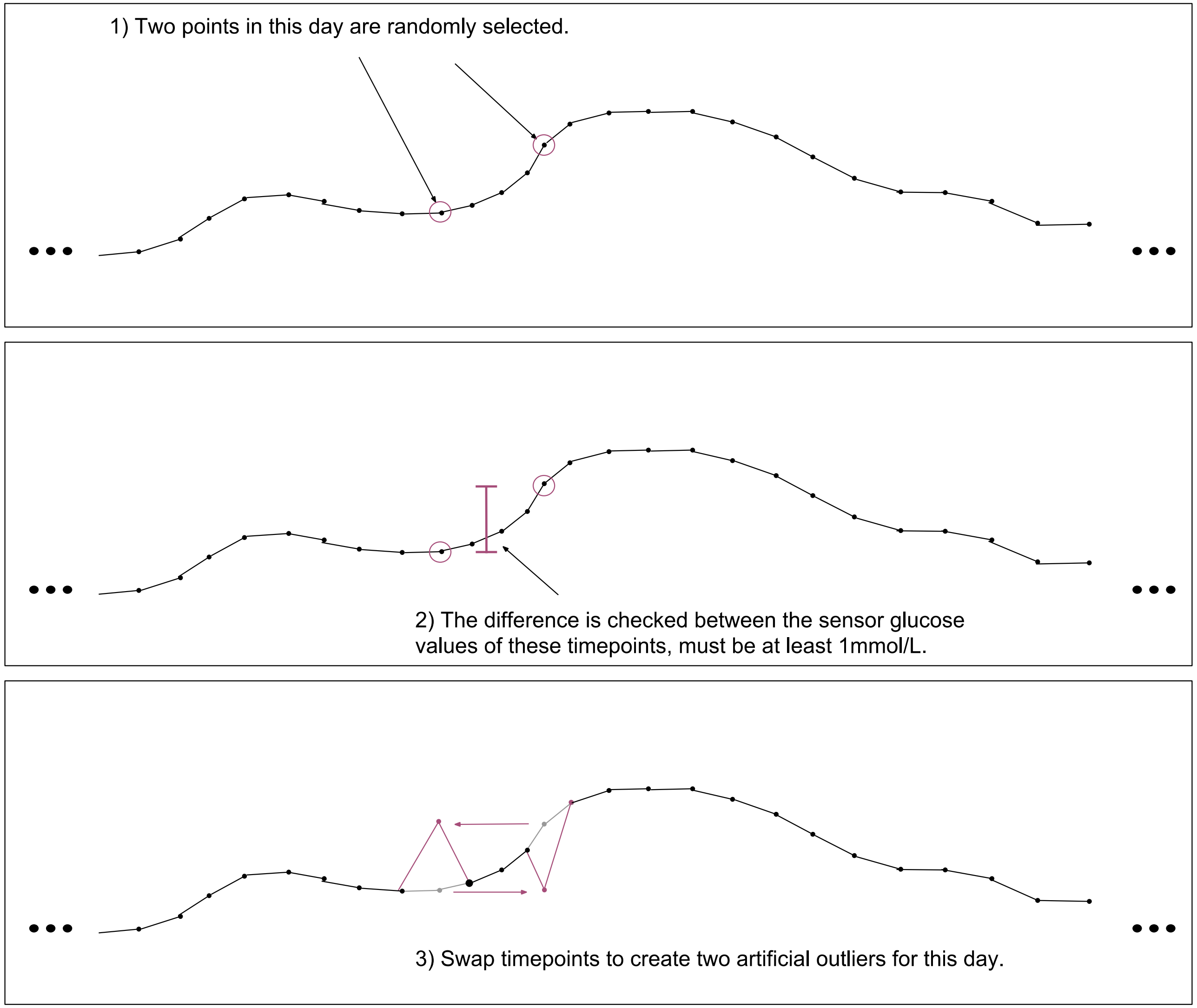
